# Towards the application of Tc toxins as a universal protein translocation system

**DOI:** 10.1101/706333

**Authors:** Daniel Roderer, Evelyn Schubert, Oleg Sitsel, Stefan Raunser

## Abstract

Tc toxins are large bacterial protein complexes that inject cytotoxic enzymes into target cells using a sophisticated syringe-like mechanism. Tc toxins are composed of a membrane translocator and a cocoon that encapsulates a toxic enzyme. The toxic enzyme varies between Tc toxins from different species and is not conserved. Here, we investigated whether the toxic enzyme can be replaced by other small proteins of different origin and properties, namely human Cdc42, herpes simplex virus ICP47, *Arabidopsis thaliana* iLOV, *Escherichia coli* DHFR, human Ras-binding domain of CRAF kinase, and tobacco etch virus protease. Using a combination of electron microscopy, X-ray crystallography and *in vitro* translocation assays, we demonstrate that it is possible to turn Tc toxins into customizable molecular syringes for delivering proteins of interest across membranes. We also infer the guidelines that protein cargos must obey in terms of size, charge, and fold in order to successfully take advantage of this new universal protein translocation system.

## Introduction

Bacteria produce and secrete an arsenal of pore-forming toxins that function by puncturing the plasma membrane of target cells. There, they either form a perforating pore that dissipates crucial electrochemical gradients or function as an injection device that translocates and releases a toxic molecule into the cytoplasm. Tripartite toxin complexes (Tc) belong to the latter class and are widespread in insect and human pathogens [1, 2]. They were originally discovered in the insect pathogen *Photorhabdus luminescens* [3] and many gene loci encoding these proteins have since been found. Tc toxins appear to be particularly well represented in enterobacteria, with prominent examples being the insect pathogen *Xenorhabdus nematophila* [4, 5], the facultative human pathogen *Photorhabdus asymbiotica* [6], and the deadly human and insect pathogens *Yersinia spp.* [7, 8].

Tc toxins consist of three components known as TcA, TcB and TcC. The ∼1.4 MDa TcA is a homopentameric bell-shaped molecule that mediates target cell association, membrane penetration and toxin translocation [9]. TcA consists of a central, pre-formed α-helical channel connected to an enclosing outer shell by a linker that acts as an entropic spring during toxin injection [10, 11]. The shell is composed of a structurally conserved α-helical domain that is decorated by a neuraminidase-like domain, as well as by highly variable immunoglobulin-fold receptor-binding domains. In some Tc toxins, the latter are functionally replaced by small soluble proteins that form a quaternary complex with the TcA subunit [12, 13].

TcB and TcC together form a ∼250 kDa cocoon that encapsulates the autoproteolytically cleaved ∼30 kDa C-terminal hypervariable region (HVR) of TcC, the actual cytotoxic component of the Tc toxin complex [10, 14]. The HVR resides in a partially or completely unfolded state in the cocoon [10, 15]. Binding of TcB-TcC to TcA and the subsequent pH-dependent prepore-to-pore transition of the ABC holotoxin result in a continuous translocation channel from the TcB-TcC lumen across the target cell plasma membrane into the cytoplasm that allows the translocation of the HVR [16]. In our previous work, we resolved several crucial steps of the Tc intoxication mechanism (reviewed in [17]). The first of these is holotoxin formation, where the key feature is a conformational transition of the TcB domain that binds to TcA. This domain is a six-bladed β-propeller, and upon contact of TcA with TcB, the closed blades of the β-propeller unfold and refold in a pseudo-symmetric open form. As a consequence, the HVR passes through the β-propeller and enters the translocation channel [16]. The assembled holotoxin binds to receptors on the target cell surface and is endocytosed [10, 18]. Upon acidification of the late endosome, the bottom of the TcA shell opens and the prepore-to-pore transition of the Tc toxin occurs [9]. During this process, the compaction of the stretched linker between the shell and the channel drives the channel through the now open bottom of the shell and across the membrane [11]. The α-helical domain of the outer shell, which possesses a stabilizing protein knot, functions as a stator for the transition [19] Subsequently the tip of the channel opens and the HVR is translocated into the target cell cytoplasm, where it interferes with critical cellular processes, ultimately causing cell death [18].

Two HVRs from *Photorhabdus luminescens* TcC proteins have been found to function as ADP-ribosyltransferases targeting actin (TccC3HVR) and Rho GTPases such as RhoA and Cdc42 (TccC5HVR) [18]. However, no HVR structures have been solved so far, limiting our understanding of the structural requirements for proteins translocated by Tc toxins. While previous studies on other bacterial ADP-ribosyltransferases have shown that these enzymes can in fact be structurally similar even without any significant sequence similarity [20–23], we do not know if this also holds true for Tc toxin HVRs.

In this study we raise an interesting question related to this topic: can the sophisticated Tc toxin translocation system be hijacked and used to transport proteins other than the natural HVRs? Such a proof of concept has already been demonstrated for the anthrax toxin, which was used to transport proteins fused to anthrax lethal factor into cells, including the TccC3HVR of Tc toxins [24]. In fact, similar designs based on the diphtheria toxin and *Pseudomonas aeruginosa* exotoxin A have already been explored as anticancer drugs [25, 26]. These systems however have the disadvantage that the fused cargo is exposed to the external environment during delivery, potentially causing the cargo to show premature and unspecific activity and limiting the usefulness of such constructs for both medical and research applications. This drawback could be avoided if the cargo were to be transported to its destination in an inactive form inside the TcB-TcC cocoon, and only activated after protein translocation through the TcA pentamer.

To achieve this, the cargo protein to be translocated would have to be fused to the C-terminus of TcC instead of the native HVR (Figure 1A,B). Naturally, it would also need to fit into the cocoon, imposing a yet to be determined upper size limit. Finally, it would also have to behave similarly enough to the HVR in terms of unfolding in the cocoon, charge and hydrophobicity distribution, for the Tc toxin to be tricked into translocating it. We explored these aspects by swapping the TccC3HVR to comparably sized proteins that display a diversity of origins and functions (Supplementary figure 1), and then assessed whether these constructs correctly form a holotoxin and translocate their cargos. We found that no stable ABC holotoxin is formed for cargos below a total size of ∼20 kDa, which is in accordance with our previous finding that an empty TcB-TcC cocoon does not form ABC with high affinity either [16]. We then screened different cargos for their translocation after triggering the prepore-to-pore transition *in vitro*. Several small proteins were successfully translocated when fused with parts of TccC3HVR. These included the intrinsically disordered infected cell protein 47 (herpes simplex virus ICP47) [27], the light / oxygen / voltage-sensing domain of the plant blue light receptor (*Arabidopsis thaliana* iLOV) [28], and the Ras-binding domain of CRAF kinase (*Homo sapiens* RBD) [29]. In contrast, we did not observe translocation of tobacco etch virus nuclear-inclusion-a endopeptidase (TEV protease) [30], dihydrofolate reductase (*Escherichia coli* DHFR) [31], and cell division control protein 42 homolog (*Homo sapiens* Cdc42) [32], with the latter two also tested as various fusions with TccC3HVR. Generally, the non-translocated cargos have a considerably lower isoelectric point (pI) than the successfully translocated constructs, with one of the Cdc42 fusion constructs and TEV being exceptions. To clarify why such cargos are not translocated, we solved the structure of TcB-TcC-Cdc42 alone and in the context of the ABC holotoxin using X-ray crystallography and electron cryomicroscopy (cryo-EM), respectively. We found that the C-terminus of Cdc42 forms an α-helix that is attached to a hydrophobic pocket inside the cocoon, where it also remains upon ABC-Cdc42 holotoxin formation, resulting in Cdc42 translocation arrest.

**Figure 1.**
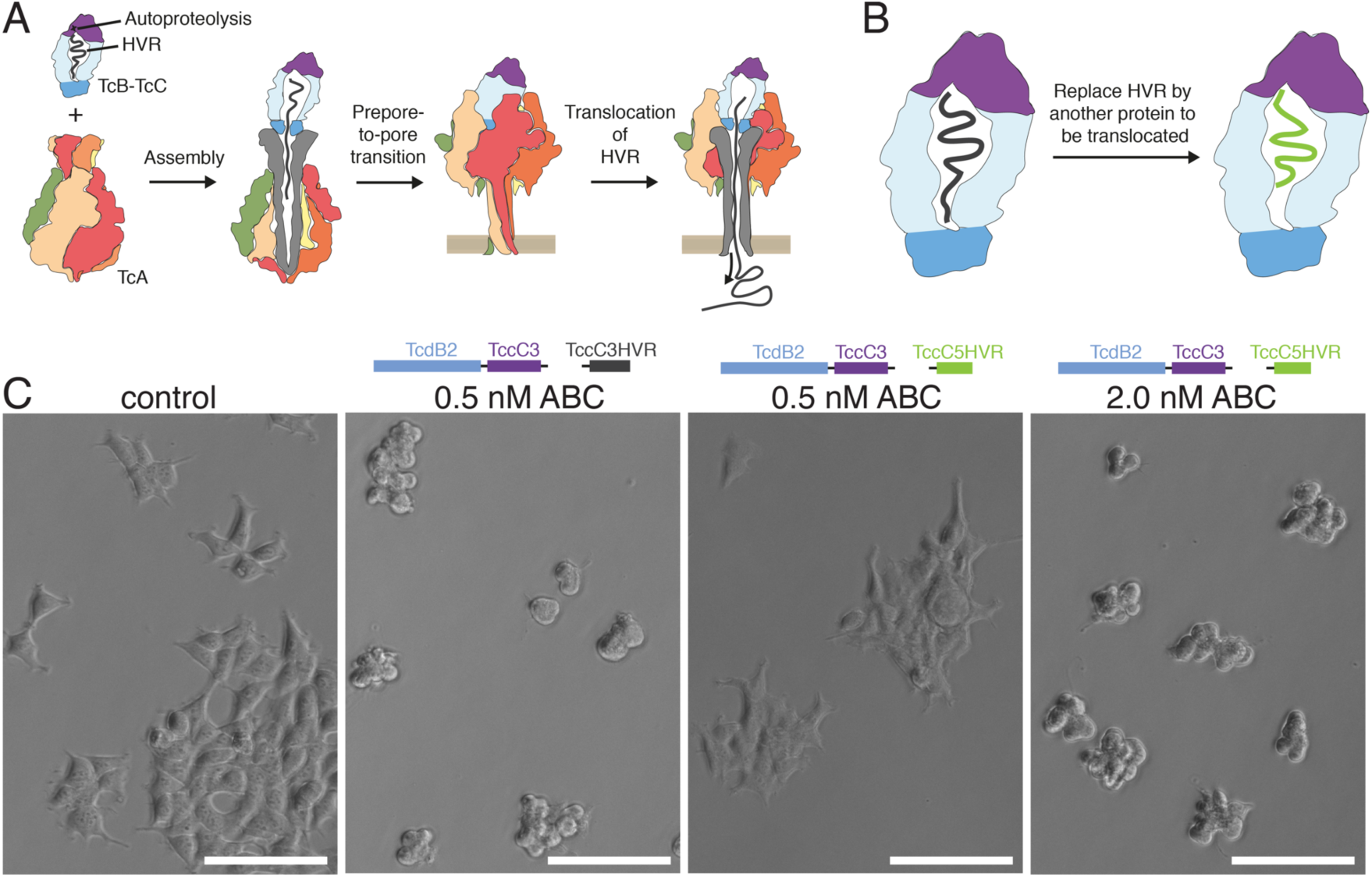
Tc toxin translocation mechanism and exchange of TccC HVRs. A. Schematic of the Tc toxin translocation mechanism. The cocoon-like TcB-TcC component (blue-purple), encapsulates the autoproteolytically cleaved cytotoxic C-terminus of TcC known as the HVR (black). Upon binding of the TcB-TcC component to the pentameric TcA component (visible monomers in red, beige, orange and green) via the TcA-binding domain of TcB (darker blue), the HVR is released into the central channel of the toxin (grey). Upon pH-induced prepore-to-pore transition at the cell membrane, the channel opens and the HVR is translocated into the cytoplasm.
B. Schematic of the experimental concept. The HVR in the cocoon is recombinantly replaced by an alternative protein to be translocated (green).
C. Effect of TccC3HVR to TccC5HVR replacement in the TcdB2-TccC3 cocoon on cytotoxicity. The ability of the TccC5HVR construct to kill HEK293T cells demonstrates that it is able to effectively translocate through the pore formed by the TcdA1 pentamer. A four-fold higher concentration of the TccC5HVR construct is needed to obtain a cytotoxic effect comparable to that of the original TccC3HVR. While this may indicate less efficient translocation, the effect is more likely due to TccC5HVR being a less potent toxin than TccC3HVR, a finding confirmed by previous studies [18]. Scale bar: 100 μm.

Together, our results show that cargo proteins must fulfill three prerequisites to be successfully translocated by TcA. The first is the cargo size, which needs to be above a threshold of about 20 kDa to form a stable holotoxin complex. The second is the net charge, which needs to be positive at neutral pH values. The third is that the cargo must not form structural elements within TcB-TcC, in particular those that interact stably with the inner surface of the cocoon. Observing these guidelines is the key to creating functional Tc-based protein injection devices.

## Results

### TcC-HVRs are exchangeable in Tc toxins

As an initial proof that different cargos fused to TcC can be translocated, we tested whether HVRs from different TcC proteins are exchangeable and result in functional, toxic ABC complexes. For this, we replaced the TccC3HVR sequence after the autoproteolytic cleavage site in TcdB2-TccC3 to that of TccC5HVR, resulting in the chimeric TcdB2-TccC3-TccC5HVR complex. After assembly of the ABC-TccC5HVR holotoxin, we assessed cytotoxicity on HEK293T cells. Complete cell death occurs upon addition of 2 nM ABC-TccC5HVR toxin, compared to cell death at 0.5 nM when exposed to the ABC-TccC3HVR holotoxin (Figure 1C). This is in accordance with previous findings that the cytotoxic effect of TccC5HVR is less pronounced than that of TccC3HVR [18]. This experiment shows that the TcdB2-TccC3 fusion protein is capable of also successfully translocating other HVRs such as TccC5HVR. We therefore chose TcdB2-TccC3 to function as a cocoon scaffold for translocation of other cargo proteins which are not components of the Tc toxin system.

### Replacement of TcC HVR by heterologous cargo proteins

Our next step was to replace the TccC3HVR with unrelated heterologous proteins and test the capability of the holotoxin to translocate these. The criteria used to select replacement proteins were i) a small size (11 – 34 kDa) to guarantee that they fit into the TcB-TcC cocoon, ii) diverse folds to assess whether this influences the translocation capability, iii) different oligomeric arrangements to see if this affects proper cocoon assembly, iv) and various organismal origins to further reduce bias. The proteins selected according to these criteria were the small GTPase Cdc42, the herpesviral ICP47 protein, the small fluorescent iLOV domain from the plant blue light receptor, the multi-ligand binding enzyme dihydrofolate reductase (DHFR), the Ras-binding domain of CRAF kinase (RBD), and tobacco etch virus (TEV) protease. Several of these proteins possess interesting properties that were hypothesized to provide additional information on requirements for translocation: Cdc42 is a homodimer in solution [33], ICP47 is intrinsically disordered, iLOV has a flavin mononucleotide chromophore, and TEV contains two β-barrels which represent stable folds (Supplementary figure 1) [34].

Interestingly, all tested chimeric cocoons could be well expressed in *E. coli* and purified. We then mixed the cocoons with TcA and assessed holotoxin formation by negative stain EM after size exclusion chromatography (SEC). In case of the wild-type TccC3HVR, the affinity of TcB-TcC to TcA is in the picomolar range, resulting in almost complete holotoxin formation [16]. This was also the case for holotoxins composed of TcA and cocoons containing Cdc42 (20.3 kDa) and TEV (28.1 kDa) (Figure 2A, Supplementary figure 2A). This demonstrates that TcB-TcC complexes with non-native cargos are able to form holotoxin complexes. In contrast, holotoxin formation was tremendously reduced when cocoons with ICP47 (11.3 kDa), iLOV (13.2 kDa), and DHFR (18.4 kDa) as cargos were used, indicating that there is a size limit for the cargo (Figure 2B, Supplementary figure 2A). Previously, we demonstrated that the HVR inside the cocoon has an influence on the gatekeeper domain of the β-propeller and an empty cocoon has a much lower affinity to TcA than the wild-type [16]. We have proposed that this might be caused by steric pressure applied by the HVR and consequently no or a small HVR would result in reduced complex formation. Our results with the different cargos support this hypothesis and indicate that the minimal size requirement for the cargo is around 20 kDa in order to guarantee high-affinity holotoxin assembly.

**Figure 2.**
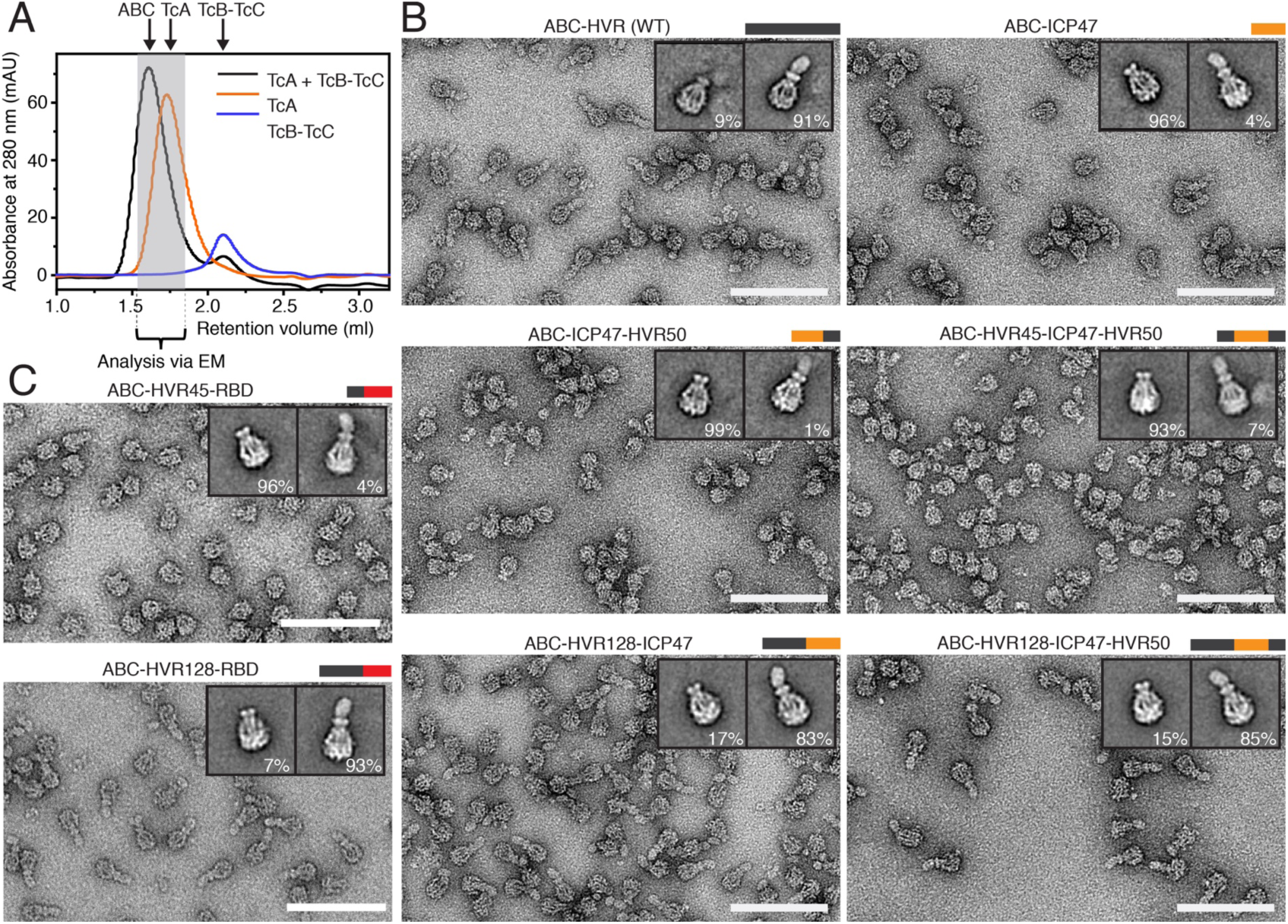
ABC holotoxin formation requires a cargo size above a distinct threshold. A. Comparison of analytical size exclusion chromatography profiles for ABC holotoxin formed by mixing TcA (600 nM) with TcB-TcC (1.2 μM), separate TcA (600 nM), and separate TcB-TcC(WT) (600 nM). The fractions indicated by the gray bar were pooled and analyzed by EM.
B. Negative stain electron micrographs of holotoxins formed by TcA and TcB-TcC with the indicated cargos (WT and ICP47 constructs). Insets: 2D class averages of representative TcA pentamers and holotoxins, with the percentages of particles in the class averages shown below.
C. Negative stain micrographs of holotoxins formed by TcA and TcB-TcC with RBD cargos, with 2D class averages like those described in panel B. While the cargo HVR45-RBD-HVR50 does not trigger holotoxin formation, the larger HVR130-RBD results in assembled holotoxins. Scale bars in B and C: 100 nm.

### The size of the cargo inside TcB-TcC determines holotoxin formation

To further explore the influence of cargo size on holotoxin formation, we created a truncated version of the native TccC3HVR (residues 1-132) and increased the size of the shorter cargos by adding differently sized parts of TccC3HVR to the N- or C-termini (Methods, Table 1). Since ICP47 has almost no secondary structure that could influence the size dependency (Supplementary figure 1), it was a particularly compelling test case. Similar to the cocoon with only ICP47 (11.3 kDa), cocoons containing the chimeras ICP47-HVR50 (17.1 kDa) or HVR45-ICP47-HVR50 (21.6 kDa) did not have a high affinity to TcA. The same was true for the short TccC3HVR (13.8 kDa) and HVR45-RBD (14.2 kDa). However, the longer constructs, namely HVR128-RBD (22.4 kDa), HVR128-ICP47 (24.5 kDa) and HVR128-ICP47-HVR50 (30.3 kDa), resulted in high-affinity holotoxin formation (Figure 2A-C). Increasing the size of iLOV (HVR128-iLOV (26.4 kDa)) and DHFR (HVR128-DHFR (31.6 kDa)) had the same effect (Supplementary figure 2A).

**Table 1.**
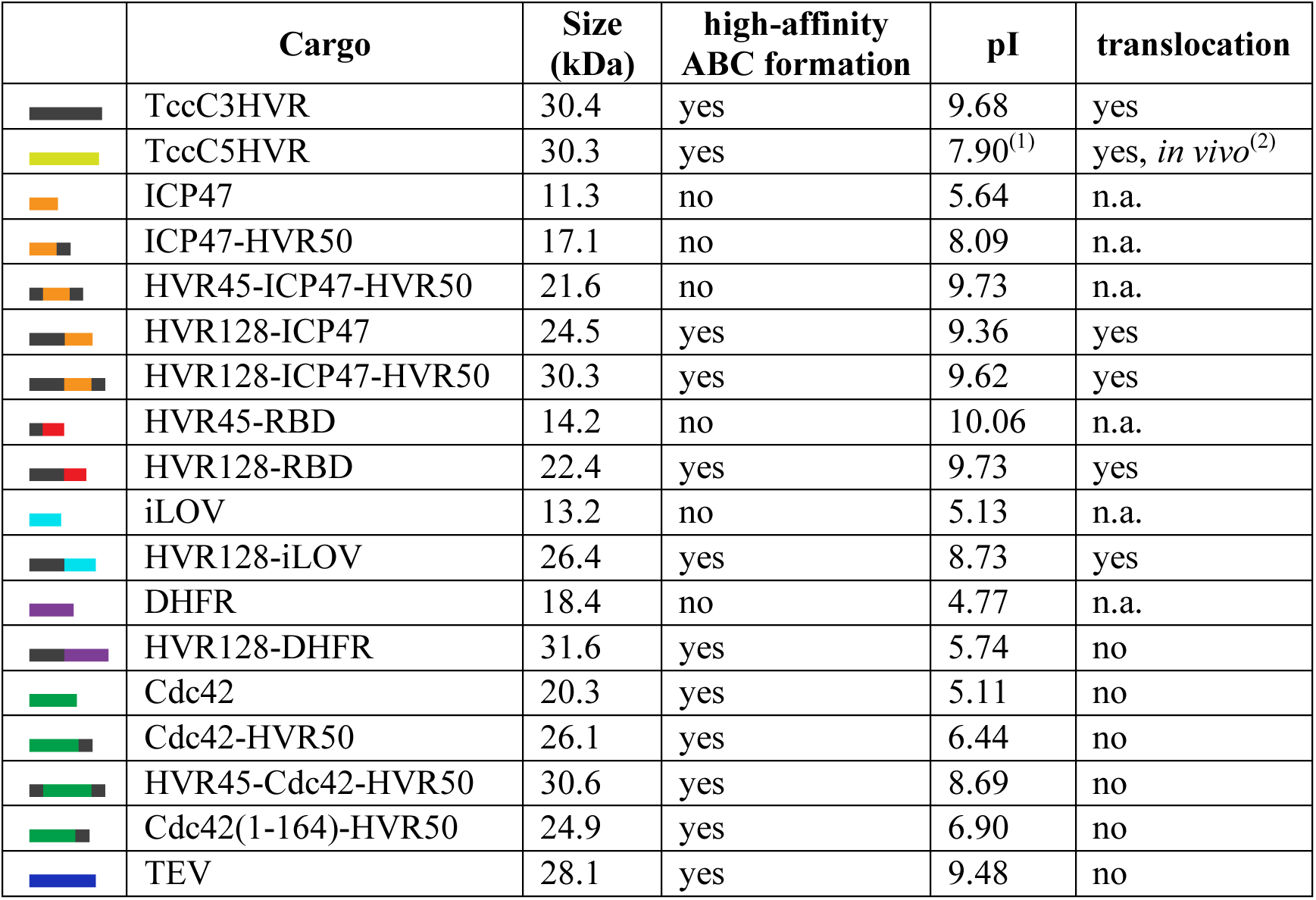
List of cargos tested in this study, including size, pI, propensity to form holotoxin, and ability to be translocated through TcA. The nomenclature of the chimeras, for example HVR128-ICP47-HVR50, indicates how many N- or C-terminal TccC3HVR residues have been pre- or appended to the cargo protein. The colored bars indicate the composition of the construct and are used in all figures to make it easier for the reader. n.a: not applicable. ^(1)^pI of TccC5HVR is changed from 8.65 to 7.90 by the addition of four residues (MPEF) to the N-terminus, resulting from cloning. ^(2)^Toxicity to HEK293T cells (Figure 1 C).

|  | Cargo | Size (kDa) | high-affinity ABC formation | pI | translocation |
| --- | --- | --- | --- | --- | --- |
|  | TccC3HVR | 30.4 | yes | 9.68 | yes |
|  | TccC5HVR | 30.3 | yes | 7.90<sup>(1)</sup> | yes, *in vivo*<sup>(2)</sup> |
|  | ICP47 | 11.3 | no | 5.64 | n.a. |
|  | ICP47-HVR50 | 17.1 | no | 8.09 | n.a. |
|  | HVR45-ICP47-HVR50 | 21.6 | no | 9.73 | n.a. |
|  | HVR128-ICP47 | 24.5 | yes | 9.36 | yes |
|  | HVR128-ICP47-HVR50 | 30.3 | yes | 9.62 | yes |
|  | HVR45-RBD | 14.2 | no | 10.06 | n.a. |
|  | HVR128-RBD | 22.4 | yes | 9.73 | yes |
|  | iLOV | 13.2 | no | 5.13 | n.a. |
|  | HVR128-iLOV | 26.4 | yes | 8.73 | yes |
|  | DHFR | 18.4 | no | 4.77 | n.a. |
|  | HVR128-DHFR | 31.6 | yes | 5.74 | no |
|  | Cdc42 | 20.3 | yes | 5.11 | no |
|  | Cdc42-HVR50 | 26.1 | yes | 6.44 | no |
|  | HVR45-Cdc42-HVR50 | 30.6 | yes | 8.69 | no |
|  | Cdc42(1-164)-HVR50 | 24.9 | yes | 6.90 | no |
|  | TEV | 28.1 | yes | 9.48 | no |

In all chimeras that led to high-affinity holotoxin assembly, the cargo was fused to HVR128. Therefore, one might ask whether these first 128 residues of TccC3HVR contain a motif important for activating TcB-TcC which the holotoxin-forming Cdc42 and TEV constructs coincidentally possess. However, since HVR(1-128) alone does not activate the cocoon, as can be seen with the HVR(1-132) truncation construct of TccC3HVR (Supplementary figure 2B), we believe that the presence of HVR128 is not a prerequisite for the activation mechanism. Taken together, these results confirm our hypothesis that the cargo has to have a certain size (at least ∼20 kDa) in order to activate the cocoon and put it into an assembly-competent state. At the same time the variety of assembly-competent constructs suggests that the nature of the cargo is not important for this activation mechanism, supporting the idea that a general high steric ‘pressure’ is sufficient.

### Translocation of cargo proteins by TcA

Having demonstrated that holotoxins containing heterologous cargos can be assembled, our next step towards using the Tc scaffold as a customized protein injection system was to show that the cargo can be successfully translocated. To assess the translocation of different cargo proteins without having to rely on protein-specific enzymatic activity readouts, we developed a cell-free *in vitro* translocation assay. First, the prepore-to-pore transition of ABC is triggered by shifting the pH to 11. If the cargo can be translocated, it will be ejected through the TcA translocation channel and dissociate from the holotoxin, as described for wild-type ABC (ABC(WT)) [15]. Successfully translocated 20 – 30 kDa cargos can then be easily separated from the 1.7 MDa ABC injection machinery by size exclusion chromatography (SEC, Supplementary figure 3A), while non-translocated cargos will still be holotoxin-associated and therefore co-migrate with the 1.7 MDa peak (Supplementary figure 3B). As proof of principle, we first assessed the release of TccC3HVR in ABC(WT) before moving on to test the heterologous cargos. Indeed, after 48 h of incubation at pH 11, a substantial fraction of TccC3HVR migrates much later than the ABC peak (Figure 3A), indicating that it has been successfully released and translocated through TcA.

**Figure 3.**
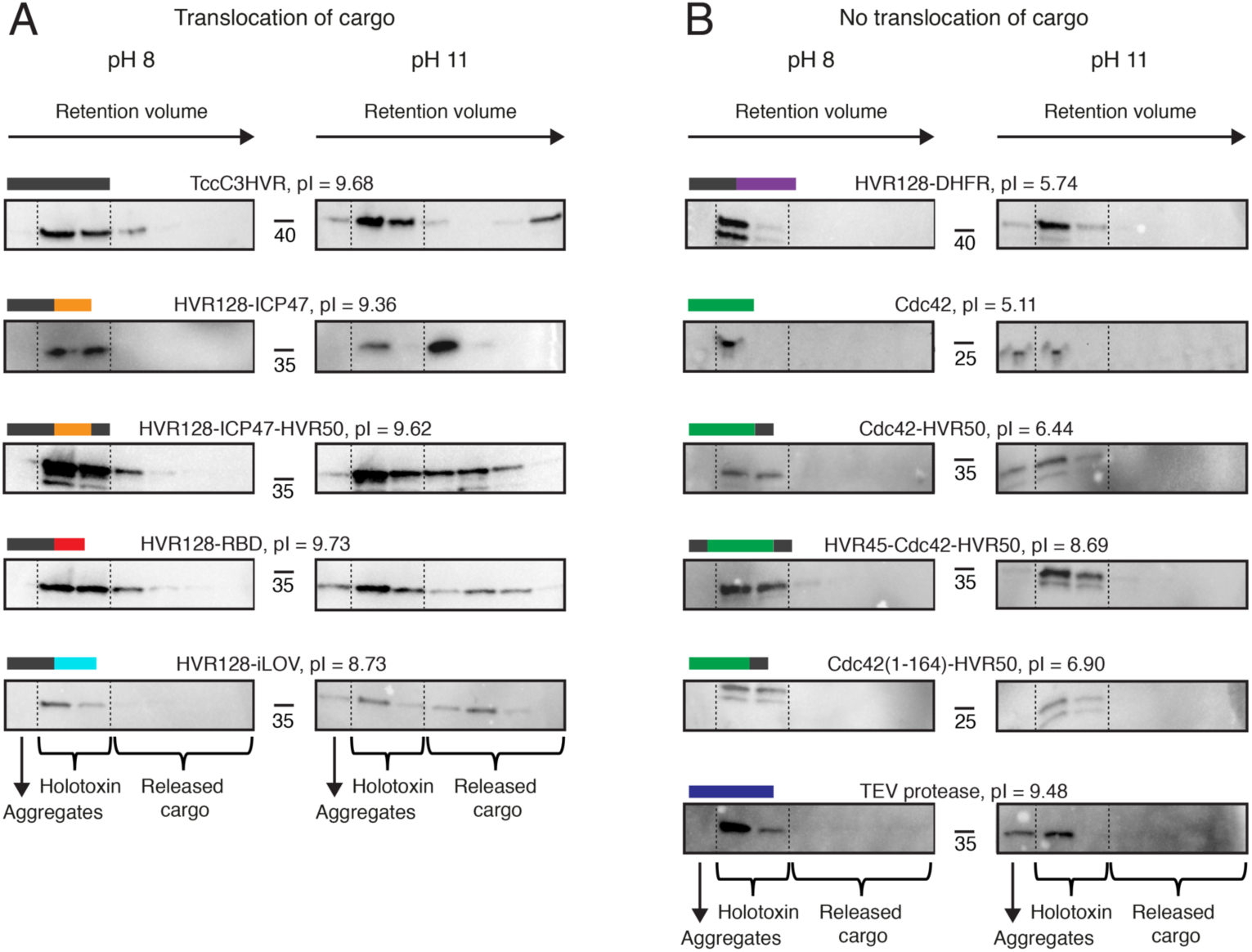
Translocation of various non-natural cargos *in vitro*. A. Successful translocation of ICP47, RBD, and iLOV fused to the N- or C-terminus of TccC3HVR. After incubation at pH 11, which causes the toxin to undergo prepore-to-pore transition, Western blots show that these constructs migrate at higher retention volumes during size exclusion chromatography in comparison to the pH 8 incubated control. This corresponds to translocation and release of the cargo proteins from the holotoxin, as illustrated schematically in Supplementary figure 3.
B. No translocation was observed for Cdc42 and TEV protease cargo alone, or DHFR and Cdc42 fused to the N- or C-terminus of TccC3HVR. Western blots show the presence of the constructs at lower retention volumes during size exclusion chromatography both at pH 8 and pH 11, meaning they co-localize with the rest of the holotoxin even after prepore-to-pore transition and are not translocated.

Next, we tested whether the various cargo proteins could be translocated. We first assessed Cdc42 and TEV, which do not need to be fused to TccC3HVR fragments in order to form a holotoxin. However, the proteins were not translocated by TcA (Figure 3B). To determine whether fusing sequences from TccC3HVR can restore translocation competence, we extended Cdc42 with either the C-terminus of TccC3HVR or with both of its termini. However, these cargos could also not be translocated by TcA (Figure 3B), indicating that neither the C-terminus nor both termini of the HVR are sufficient to enable translocation.

However, the four other cargos that facilitated holotoxin formation only when fused to fragments of TccC3HVR were successfully translocated, namely HVR128-ICP47-HVR50, HVR128-ICP47, HVR128-iLOV and HVR128-RBD (Figure 3A). Since the C-terminal region that is translocated first in ABC(WT) [16] differs considerably between these cargos, we conclude that there is no specific sequence at the C-terminus that determines whether a cargo is translocated or not. In the case of HVR128-iLOV, we did not observe iLOV fluorescence in the cocoon (Supplementary figure 3C), indicating that it is stored in an unfolded form. In contrast to the other three HVR128-containing cargos, DHFR is not translocated when fused to the same HVR128 N-terminus (Figure 3B). Together with the non-translocated HVR45-Cdc42-HVR50 cargo, this shows that the sequence of the N-terminus is also not the determinant of cargo transport through TcA. Therefore, there must be another factor at work that establishes translocation competence.

A comparison of the four translocated fusion proteins (HVR128-ICP47, HVR128-ICP47-HVR50, HVR128-RBD and HVR128-iLOV) and the native cargos Tcc3HVR and Tcc5HVR shows that their common feature is a positive net charge at neutral pH, with isoelectric points of at least 7.9 (Figure 3A). In the case of the fusion constructs, this is mainly due to the highly positively charged HVR128 (pI 9.75), which is larger than the cargo protein in all cases (maximum size 13.2 kDa). We therefore conclude that besides being large enough, the cargo has to be positively charged (pI *≥* ∼8) in order to be translocated.

In line with this, the six constructs that formed holotoxin complexes but did not show translocation had mostly negatively charged cargos (Table 1). Only two non-translocated cargos, namely HVR45-Cdc42-HVR50 and TEV were positively charged. The TEV construct we used has a highly positively charged penta-arginine tail to allow purification by cation exchange chromatography without changing its activity [35], a modification that raises the pI of TEV from 8.67 to 9.62. While such a change in pI should favor translocation, it is possible that distributing the positive charges in such an uneven manner rather hinders it. In addition, TEV contains two β-barrels (Supplementary figure 1), which are known to be a notoriously stable fold [34]. For the native Tcc3HVR we have shown that it resides in the TcB-TcC cocoon in an unfolded or partially unfolded state [15]. This is further supported by the fact that HVR128-iLOV is non-fluorescent in the cocoon (Supplementary figure 3C), indicating that it is stored in an unfolded conformation. Since folded proteins do not fit through the narrow constriction site of the translocation channel [16], the formation of defined structural elements such as β-barrels in the cocoon probably result in translocation arrest.

### The interaction of an α-helix with the cocoon inhibits translocation of Cdc42

Although Cdc42 does not possess any folds that are immediately classifiable as very stable, we looked at the Cdc42 constructs in more detail and solved the crystal structure of TcB-TcC-Cdc42 to 2.0 Å to find out whether Cdc42 folding inside the cocoon is nonetheless a plausible explanation for its translocation incompetence (Figure 4A, Supplementary Table 1).

**Figure 4.**
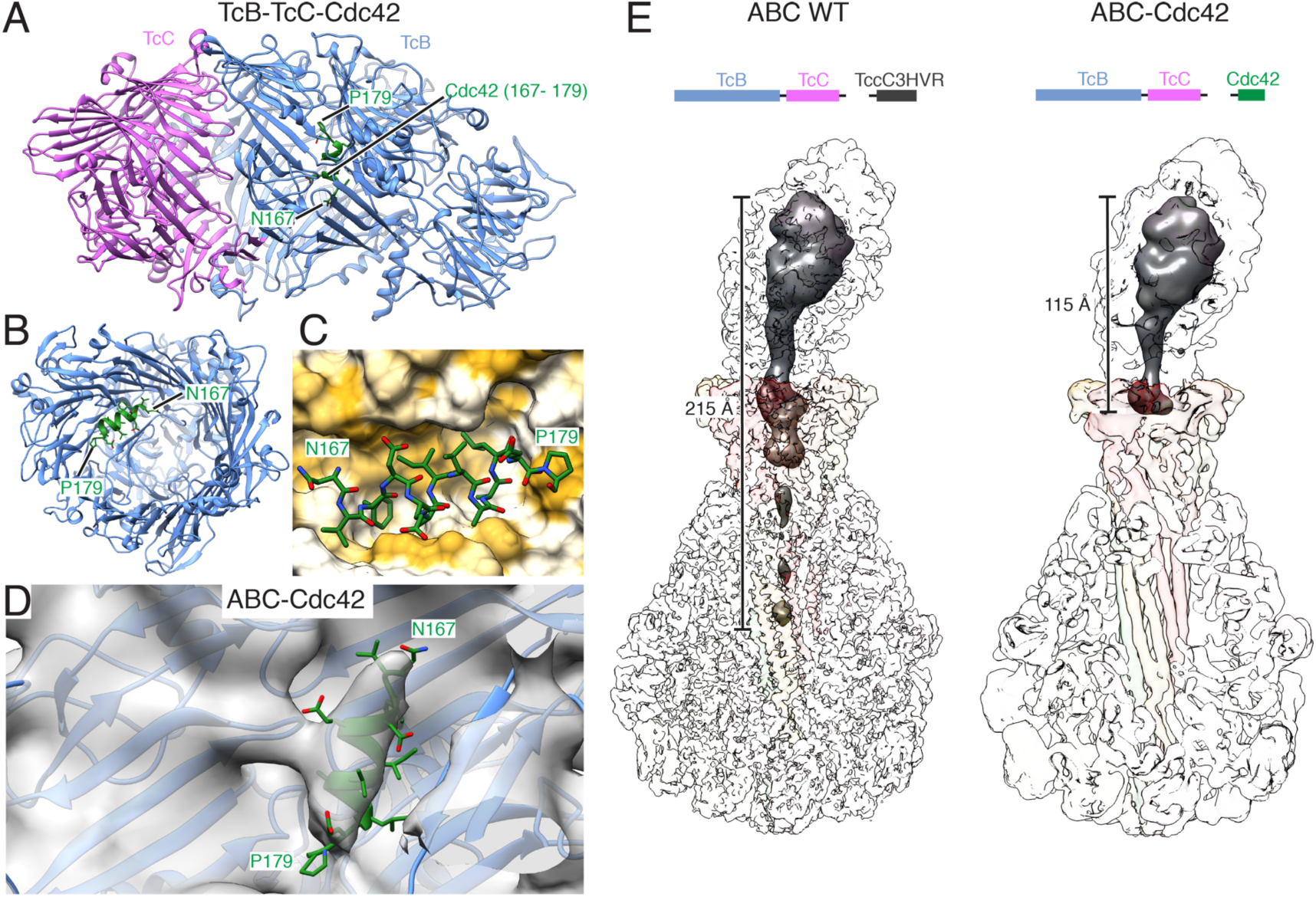
Structure of the fusion protein TcB-TcC-Cdc42 alone and in complex with TcA. A. Crystal structure of the TcB-TcC-Cdc42 complex. The C-terminal region of Cdc42 (N167 – P179, green) forms a helix inside the cocoon.
B. Top view into TcB-TcC-Cdc42, with the TcC model removed for illustrative purposes. The C-terminal Cdc42 α-helix (green) can be seen attached to one side of the cocoon, while the remaining Cdc42 density is too disordered to resolve.
C. Surface representation of the binding pocket of the hydrophobic C-terminal Cdc42 helix in the TcB section of the cocoon. The molecular surface of TcB is colored according to its hydrophobicity [47], with hydrophobic regions shown in ochre and polar regions in white.
D. Rigid-body fit of the crystal structure of TcB-TcC-Cdc42 into the cryo-EM density map of ABC-Cdc42. Density corresponding to the α-helix of Cdc42 is also present in the cryo-EM structure of the holotoxin at the same site.
E. Comparison of cryo-EM density reconstructions of ABC(WT) (PDB ID 6H6F). and ABC-Cdc42 (this work). The surfaces of the TcA, TcB, and TcC subunits are transparent, and the density corresponding to the ADP-ribosyltransferase TccC3HVR (WT, left) and Cdc42 (right) is dark grey. The latter is shown at a lower threshold and filtered to 15 Å resolution.

The overall shape of the TcB-TcC-Cdc42 cocoon is identical to wild-type and empty TcB-TcC [10, 16], indicating that a different cargo does not influence the RHS repeat structure of the cocoon (Supplementary figure 4A). Similar to the native TcB-TcC cocoon, Cdc42 inside the cocoon is not structured. However, we found an ordered electron density in close spatial proximity to the β-propeller domain, which corresponds to an α-helix not present in other TcB-TcC structures obtained so far. The high resolution of the map and availability of the Cdc42 structure [32] enabled us to identify the α-helix inside the cocoon as the C-terminus of Cdc42 (N167 – P179) (Supplementary figure 4B,C). Importantly, the helix is attached to the TcB protein in an orientation perpendicular to that expected from natural translocation (Figure 4A,B), and its amphipathic nature facilitates interaction with a hydrophobic pocket on the inner surface of the cocoon. The side chains of F169, I173, L177 and P179 are rigidly oriented towards the hydrophobic cleft, while D170, E171 and E178 face the TcB-TcC lumen with more degrees of freedom (Figure 4C). This is reflected in the quality of the crystallographic density, with only the residues pointing towards the cocoon surface being well resolved (Supplementary figure 4D). Interestingly, the affinity of the Cdc42 α-helix for this part of the cocoon is strong enough to displace the N-terminus of TcB, which resides at this position in TcB-TcC(WT) (Supplementary figure 4E). The N-terminus of TcB-TcC-Cdc42 is correspondingly not resolved, indicating that it protrudes into the cocoon lumen. The tight attachment of Cdc42 via its C-terminus in close spatial proximity to the TcB gatekeeper domain [16] could help this construct to form a holotoxin with high affinity comparable to the constructs with larger cargos, despite Cdc42 being at the lower limit of the cargo size prerequisite (Figure 2, Supplementary figure 2).

The stable nature of the Cdc42 α-helix interaction with the cocoon raises the question of whether it remains bound even after holotoxin formation, in which case it would not be able to enter the translocation channel. We addressed this issue by determining the 5 Å structure of the ABC-Cdc42 holotoxin using cryo-EM (Supplementary figure 5). Indeed, a small helix-shaped density appears at the interaction site even at a comparably high map binarization threshold, despite the limited resolution of the 3D reconstruction. Fitting the entire crystal structure of TcB-TcC-Cdc42 into the cryo-EM map results in a very good match of the Cdc42 α-helix with the additional map density of ABC-Cdc42 (Figure 4D). In contrast, no comparable cryo-EM density is present at the same position in ABC(WT) and an aspartyl-protease deficient variant [16] (Supplementary figure 6), indicating that the cargo does not form any structural elements at this location in functional holotoxins.

**Figure 5.**
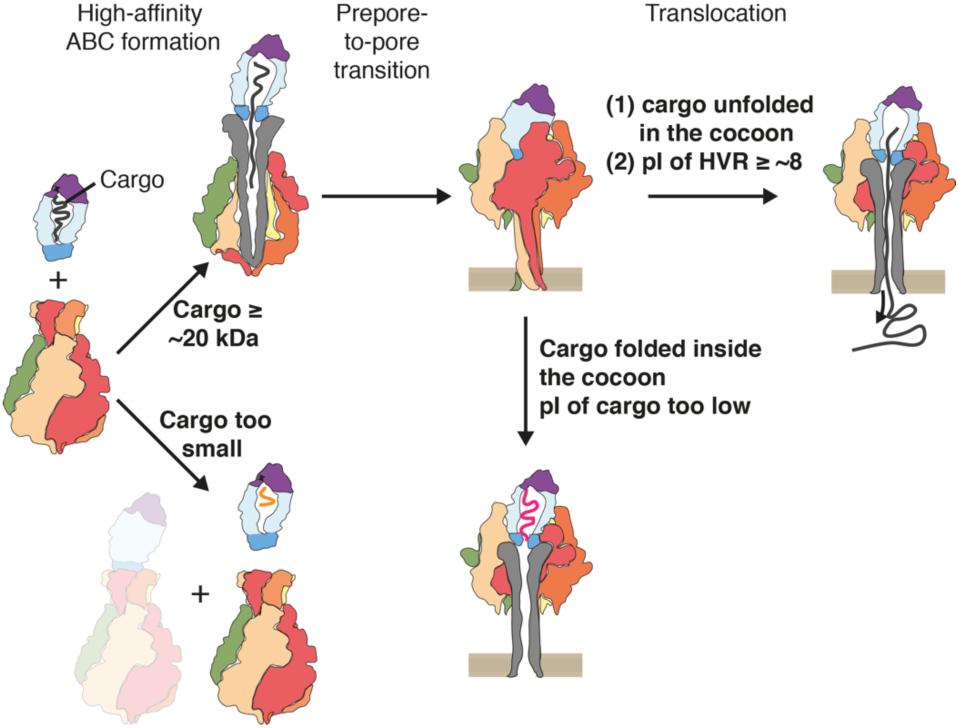
Prerequisites for creating a functional customized Tc translocation system and overview of tested cargos. TcA and TcB-TcC assemble into a holotoxin with high affinity only when the cargo is sufficiently large (*≥* 20-22 kDa). Translocation upon prepore-to-pore transition only occurs if the cargo is unfolded in the cocoon and its pI is *≥* 8.

The tight association of the C-terminal α-helix to the cocoon results in Cdc42 translocation arrest already at an early stage. If the C-terminus that would normally be translocated first through TcA [16] is occupied, then why does Cdc42 form a holotoxin? Analysis of the cryo-EM data shows that no additional density is present in the TcA translocation channel, unlike in ABC(WT) (Figure 4E). There is however a stretch of Cdc42 density that protrudes into the β-propeller domain of TcA but does not continue further into the channel. This indicates that while Cdc42 cannot be translocated, it applies a steric pressure on the TcB gatekeeper domain, resulting in the high-affinity binding of TcB to TcA.

These results demonstrate that if the encapsulated cargo forms stable structural elements that strongly interact with the inner lumen of the cocoon, the cargo cannot be translocated by TcA even if it is positively charged and large enough.

## Discussion

Taken together, our results indicate that the *P. luminescens* Tc toxin can be successfully transformed into a universal protein translocation system as long as the cargo protein fulfills several prerequisites. The first of these is cargo size. While its upper limit is defined by the size of the cocoon, an exact value was not determined in this study. The largest cargo tested here was HVR128-DHFR (31.6 kDa), which is 1.2 kDa larger than the TccC3HVR natural cargo (Table 1). At the same time, the largest natural cargo of a *P. luminescens* Tc toxin is TccC1HVR at 35.1 kDa, providing a potential upper size limit and meaning that larger ADP-ribosyltransferases like the 49.4 kDa C2 toxin of *Clostridium botulinum* [21] would likely not fit into the cocoon. It remains to be explored whether the cocoon itself can be enlarged by adding further RHS repeats, thereby expanding the upper cargo size limit.

Importantly, we discovered that the lower size limit of the cargo is in the 20 – 22 kDa range, and the affinity of TcB-TcC to TcA drops tremendously if the cargo size is below this limit (Figure 5, Table 1). This finding echoes previous results showing that a comparable decrease in affinity between empty TcB-TcC and TcA occurs because there is no HVR present to apply steric pressure on the TcB gatekeeper domain that is essential for binding to TcA [16]. Similarly, small cargos will likely be too mobile inside the cocoon to cause gatekeeper destabilization. The lower size limit is slightly flexible, e.g. Cdc42 (20.3 kDa) forms holotoxin while HVR45-ICP47-HVR50 (21.6 kDa) does not (Table 1). This is potentially caused by the C-terminal helix of Cdc42 attaching the rest of the cargo to the vicinity of the cocoon exit, allowing the Cdc42 N-terminus to destabilize the gatekeeper more easily, which results in holotoxin formation but also prevents subsequent cargo transport. We therefore hypothesize that translocation will likely be impaired if a TcB-TcC cocoon loaded with a small cargo binds with high affinity to TcA.

The second prerequisite for successful transport is a net positive charge of the cargo protein. All naturally occurring HVRs of *P. luminescens* Tc toxins have a pI of at least 7.9 (TccC5HVR), and the protein with the lowest pI for which we observed translocation in our *in vitro* assay is HVR128-iLOV with a pI = 8.73 (Table 1). Since the gatekeeper constriction site that serves as the entry point into the TcA channel is formed by several acidic residues [16], negatively charged constructs might be repelled. In addition, the translocation channel of TcA has several negatively charged bands [10, 11] that only facilitates the transport of cations and not of anions [10, 36].

A sufficiently large size and a positive net charge are however not the only points that need to be considered, because some of the larger cargos with high pI were not translocated (TEV and HVR45-Cdc42-HVR50). Therefore, a third prerequisite needs to be fulfilled: the encapsulated cargo must not form any tertiary structures or stably interact with the cocoon. The two abovementioned cargos violate this prerequisite, with TEV containing two highly stable β-barrels, and Cdc42 possessing an amphipathic C-terminus that associates with a hydrophobic binding pocket in the cocoon.

To summarize, we recommend adhering to the following guidelines in order to successfully turn a Tc toxin into a protein translocation device (Figure 5):

1. Choose a protein between 20 and 35 kDa to insert after the autoproteolytic cleavage site of TcC. In case the cargo is too small, add N- or C-terminal extensions that do not interfere with protein function or link several copies of the protein together.
2. Choose a protein with a pI of at least 8.0. Increase the pI by addition of positively charged residues or site-directed mutagenesis if necessary and possible.
3. Avoid cargo proteins that might form highly stable structures already in the cocoon lumen.
4. Avoid cargo proteins that contain amphipathic helices or extensive hydrophobic patches that might stably interact with the inner surface of the cocoon.

## Acknowledgements

We thank K. Vogel-Bachmayr for excellent technical support. We thank R. Tampé for a plasmid containing the ICP47 sequence and A. Ernst for a plasmid containing the RBD sequence. This work was supported by funding from the Max Planck Society (to S.R.) and the European Research Council under the European Union’s Seventh Framework Programme (FP7/2007-2013 grant no. 615984, to S.R.).

## Author contributions

S.R. designed and supervised the project. D.R. designed proteins. D.R. performed the assembly and translocation experiments. D.R. and O.S. performed fluorescence spectroscopy. E.S. crystallized TcB-TcC-Cdc42, and solved the crystal structure. D.R. processed and analyzed cryo-EM data of ABC-Cdc42. D.R., O.S. and S.R. wrote the manuscript.

## Author information

The coordinates for the EM structure of ABC-Cdc42 have been deposited in the Electron Microscopy Data Bank under accession number ….. The coordinates of the crystal structure of TcB-TcC-Cdc42 have been deposited in the Protein Data Bank under accession number … Correspondence and requests for materials should be addressed to S.R.

## Methods

### Source plasmids for cargo proteins

pGEX-iLOV was a gift from John Christie (Addgene plasmid #26587) [28]. pcDXc3A-AN44-ICP47 containing the coding sequence (CDS) for ICP47 with a C-terminal HA epitope was a gift from Robert Tampé, Goethe University of Frankfurt, Germany. pGEX-RBD with the CDS for the Ras-binding domain (RBD) of CRAF kinase was a gift from Andreas Ernst, Goethe University of Frankfurt, Germany. A customized plasmid with the CDS for dihydrofolate reductase (DHFR) was a gift from Yaowen Wu, Max Planck Institute of Molecular Physiology, Dortmund, Germany. pOPIN-MBP-TEV with the CDS for TEV protease with a C-terminal penta-arginine tail was a gift from the Dortmund Protein Facility (DPF), Max Planck Institute of Molecular Physiology, Dortmund, Germany.

### Cloning of TcB-TcC with non-natural cargo proteins

We used the fusion protein TcdB2-TccC3 [10] as a carrier for heterologous cargo proteins to replace TccC3HVR. We therefore introduced an EcoRI restriction site by site-directed mutagenesis after the codon coding for the conserved P680 of TccC3, which is two residues after the aspartyl protease site [15]. Subsequently, we cloned the cargo proteins in frame via EcoRI and XhoI, resulting in an N-terminal extension of four residues (MPEF). In the case of TccC5HVR, the N-terminal extension results in a change of the pI from 8.65 to 7.90. In the case of Cdc42, the EcoRI site was inserted after the codon for L678, and the resulting point mutation P680F was reverted by site-directed mutagenesis. Cargo proteins with N-terminal extensions were generated by inserting the EcoRI site 135 or 384 base pairs after the aspartyl protease site and restriction cloning as described above, resulting in 45 or 128 N-terminal residues of TccC3HVR and two residues (EF) encoded by the EcoRI site being attached to the cargo. Cargo proteins with C-terminal extensions were generated by inserting a NotI restriction site 150 base pairs upstream of the stop codon, followed by restriction cloning via EcoRI and NotI. This resulted in a C-terminal extension of the cargo proteins containing 50 C-terminal residues of TccC3HVR and three residues (GGR) encoded by the NotI site. An overview of all resulting TcB-TcC cargos is listed in Table 1.

### Protein production

*P. luminescens* TcdA1 (TcA) was expressed and purified as described previously [16]. *E. coli* BL21-CodonPlus(DE3)-RIPL cells were transformed with pET19b containing the *tcdA1* gene with an N-terminal hexahistidine tag and a pre-culture was inoculated from a freshly transformed colony. 10 L LB medium were inoculated to an OD600 of 0.05 and incubated at 37 °C. At an OD600 of 0.6, expression was induced by 25 μM IPTG and carried out overnight at 20 °C. Cells were disrupted in lysis buffer (20 mM Tris-HCl pH 8.0, 200 mM NaCl, 5 mM imidazole, 0.05% Tween-20) using a microfluidizer. After removal of cell debris by centrifugation, the supernatant was applied to a 5 mL Ni-NTA column (GE Healthcare Life Sciences) and washed with 10 column volumes (CVs) of washing buffer (lysis buffer with 50 mM imidazole). Subsequently, the protein was eluted with elution buffer (lysis buffer with 150 mM imidazole). Eluted TcdA1 was dialyzed against 20 mM HEPES-NaOH pH 8.0, 150 mM NaCl, 0.05% Tween-20 and size exclusion chromatography (SEC) on a Sephacryl S400 16-60 column (GE Healthcare Life Sciences) equilibrated in the same buffer was performed as last purification step.

All TcB-TcC cargo variants were expressed and purified analogously to WT TcB-TcC as described previously [11]. *E. coli* BL21-CodonPlus(DE3)-RIPL cells were transformed with pET28a encoding the *tcdB2-tccC3* (WT or cargo variants) genes with an N-terminal hexahistidine tag. 5 or 10 L of LB medium containing 30 mM IPTG were directly inoculated with a freshly transformed colony. Cells were grown at 28 °C for 4 h, followed by 25 °C for 20 h and 20 °C for 24 h. Subsequently, cells were disrupted in lysis buffer (20 mM Tris-HCl pH 8.0, 300 mM NaCl, 10 % glycerol) using a microfluidizer. After removal of cell debris, the supernatant was applied to a 5 mL Ni-NTA column (GE Healthcare Life Sciences) and washed with 10 CVs of washing buffer (lysis buffer with 40 mM imidazole), followed by elution with a linear gradient from 40 mM to 250 mM imidazole over 10 CVs. The eluted protein was diluted with dilution buffer (20 mM Tris-HCl pH 8.0, 5% glycerol) to a final NaCl concentration of 20 mM and immediately loaded on a 5 mL HiTrapQ column (GE Healthcare Life Sciences) as a second purification step. After washing with 10 CVs of washing buffer 2 (20 mM Tris-HCl pH 8.0, 20 mM NaCl, 5% glycerol), the protein was eluted with a linear gradient from 20 to 500 mM NaCl over 20 CVs. Fractions containing TcB-TcC were subjected to SEC using a Superdex 200 10-300 or a Superdex 200 16-60 column (GE Healthcare Life Sciences) equilibrated in gel filtration buffer (20 mM Tris-HCl pH 8.0, 150 mM NaCl, 5% glycerol) as final purification step.

### ABC holotoxin formation

Purified TcA (600 nM pentamer) and different TcB-TcC variants (1.2 μM) were mixed together and incubated for at least 1 h at 4 °C before removing the excess of free TcB-TcC by SEC on a Superose 6 Increase 5-150 column (GE Healthcare Life Sciences). For TcB-TcC variants containing HVR-RBD chimeras, 300 nM of TcA pentamer and 600 nM of TcB-TcC were used instead. The formation of the resulting ABC variants was verified by negative stain electron microscopy.

### Negative stain electron microscopy

After SEC, 3 μL of 0.1 mg/mL of each ABC variant were incubated for 1 min on a glow-discharged 400-mesh copper grid (Agar Scientific) with an additional layer of thin carbon film. Subsequently, the sample was blotted with Whatman no. 4 filter paper and stained with 0.75% uranyl formate. Images were recorded on a FEI Tecnai Spirit TEM operating at 120 kV and equipped with a F416 CMOS detector (TVIPS). To quantify the ratio of holotoxin assembly, we picked at least 1000 particles from every ABC variant using crYOLO [37] and subjected them to 2D classification with ISAC [38] in SPHIRE [39].

### Cell intoxication

HEK 293T cells were intoxicated with WT ABC or ABC-TccC5HVR. Cells (5 × 10^4^ per well) were grown adherently overnight in 400 µL DMEM/F12 medium (Pan Biotech) and 0.5 or 2 nM holotoxin was subsequently added. Incubation was performed for 16 h at 37 °C before imaging. Experiments were performed in triplicate. Cells were not tested for *Mycoplasma* contamination.

### *In vitro* protein translocation assay

After ABC formation and removal of unbound TcB-TcC via SEC, 200 nM ABC (WT or cargo variants) was mixed with DDM (final concentration: 0.1%) and dialyzed against 20 mM CAPS-NaOH pH 11.2, 150 mM NaCl, 0.1% DDM for 48 h at 4 °C. As a parallel control experiment, the same amount of ABC with 0.1% DDM was dialyzed against 20 mM Tris-HCl pH 8.0, 150 mM NaCl, 0.1% DDM under the same conditions. Subsequently, the dialyzed proteins were subjected to SEC on a Superose 6 Increase 5-150 column equilibrated in the respective dialysis buffer. SEC fractions corresponding to the exclusion volume (aggregated holotoxin after dialysis), the major peak of the holotoxin, the tail of the holotoxin peak, and the peak after holotoxin were analyzed via SDS-PAGE and Western blot for the presence of the cargo. In the cases where the cargo is translocated and released from the holotoxin, a Western blot signal appears in the fractions after the holotoxin peak at pH 11 (Supplementary figure 3A). In the cases where the cargo is not translocated, Western blot signals are only found in fractions containing the holotoxin, both at pH 8 and pH 11 (Supplementary figure 3B).

### Western blot and immunodetection

After SDS-PAGE of the collected SEC fractions (10 μL per fraction) on a 4-15% acrylamide gradient gel, the proteins were transferred onto a PVDF membrane using a Trans-Blot Turbo semi-dry transfer system (Biorad). In the cases where the cargo proteins were fusion constructs with N- or C-terminal parts of TccC3HVR, a custom-made anti-TccC3HVR rabbit polyclonal antibody (Cambridge Research Biochemicals) was used as the primary antibody. For Cdc42 without a TccC3HVR fusion, an anti-Cdc42 rabbit polyclonal antibody (Cell Signaling Technology, Cat. No. 2462) was used as the primary antibody. For TEV, an anti-TEV protease rabbit polyclonal antibody (Novus Biologicals, Cat. No. NBP1-97669) was used as the primary antibody. An HRP-conjugated goat anti-rabbit antibody (Biorad, Cat. No. 170-6515) was applied as the secondary antibody in all cases. Detection was performed with Western Lightning Plus ECL reagent (PerkinElmer, Cat. No. NEL104001EA) and imaged in a ChemiDoc MP imaging system (Biorad).

### Fluorescence spectroscopy

Florescence emission spectra of TcB-TcC(WT), TcB-TcC-iLOV and TcB-TcC-HVR130-iLOV (500 nM protein concentration) were recorded in 20 mM Tris-HCl pH 8.0, 150 mM NaCl, 0.05% Tween-20 using a FluoroMax-4 fluorescence spectrophotometer (Horiba). The excitation wavelength was 450 nm, and emission was recorded from 490 to 620 nm. Purified iLOV with a GST-tag [28] was used as positive control.

### X-ray crystallography of TcB-TcC-Cdc42

TcB-TcC-Cdc42 was crystallized using the sitting-drop vapor diffusion method at 20 °C. Initially, 2D crystals formed after mixing 1 µL of 10 mg/mL protein solution with 1 µL reservoir solution containing 0.1 M sodium chloride, 0.1 M magnesium chloride, 0.1 M tri-sodium citrate pH 5.5 and 12 % PEG 4000. The 2D crystals were used to prepare a seed solution. Final 3D crystals were obtained by mixing 1 µL of 10 mg/mL protein solution with 0.5 µL seed solution and 1.5 µL reservoir solution containing 0.1 M magnesium chloride, 0.1 M tri-sodium acetate pH 4.6 and 12 % PEG 6000. Prior to flash freezing in liquid nitrogen, the crystals were soaked in reservoir solution containing 20 % glycerol as a cryoprotectant.

X-ray diffraction data was collected at the PXII-X10SA beamline at the Swiss Light Source (Villigen, Switzerland) using a wavelength of 0.97958 Å. The X-ray dataset was integrated and scaled using XDS [40]. Phases were determined by molecular replacement with PHASER implemented in PHENIX [41] using the crystal structure of WT TcdB2-TccC3 (PDB ID 4O9X) as a search model. TcB-TcC-Cdc42 crystallized in the orthorhombic space group P212121 with unit cell dimensions of 96 × 156 × 179 Å and one molecule per asymmetric unit. The structures were optimized by iteration of manual and automatic refinement using COOT [42] and PHENIX [43] to a final Rfree of 25%. Data collection and refinement statistics are summarized in Supplementary Table 1.

### Cryo-EM sample preparation and data acquisition of ABC-Cdc42

3 µL of 2.1 mg/mL ABC-Cdc42 in 20 mM Tris-HCl pH 8.0, 150 mM NaCl, 0.05% Tween-20 were applied on a glow-discharged holey carbon grid (Quantifoil, QF 2/1, 300 mesh). Subsequently, the sample was vitrified in liquid ethane with a Cryoplunge3 plunger (Cp3, Gatan) using 1.6 s blotting time at 90 % humidity and 22 °C.

A dataset of ABC-Cdc42 was collected at the Max Planck Institute of Molecular Physiology, Dortmund using a Cs corrected Titan Krios equipped with an XFEG and a Falcon II direct electron detector. Images were recorded using the automated acquisition program EPU (FEI) at a magnification of 59,000x, corresponding to a pixel size of 1.14 Å/pixel on the specimen level. 3024 movie-mode images were acquired in a defocus range of 1.0 to 3.2 µm. Each movie comprised 24 frames with a total cumulative dose of **∼**65 e^-^/Å^2^.

### Image processing of ABC-Cdc42

After initial screening of all micrographs, 2754 images were selected for further processing. Movie frames were aligned, dose-corrected and averaged using MotionCor2 [44]. The integrated images were also used to determine the contrast transfer function (CTF) parameters with CTER [45], implemented in the SPHIRE software package [39]. Initially, 2025 particles were manually picked and 2D class averages generated by Relion [46] were used as an autopicking template. 99,980 particles were auto-picked from the images using the Relion 1.4 autopicker. Subsequently, reference-free 2D classification and cleaning of the dataset were performed with the iterative stable alignment and clustering approach ISAC [38] in SPHIRE. ISAC was executed with a pixel size of 7.2 Å/pixel on the particle level. The ‘Beautifier’ tool of SPHIRE was then applied to obtain refined and sharpened 2D class averages at the original pixel size, showing high-resolution features (Supplementary figure 5b). From the initial set of particles, the clean set used for 3D refinement contained 56,665 particles. We applied the previously obtained cryo-EM structure of ABC(WT) (EMDB-2551) as an initial model after scaling and filtering it to 12 Å resolution and performed 3D refinement in SPHIRE. The resolution of the final density was estimated to be 7.02 / 5.11 Å according to FSC 0.5 / 0.143 after applying a soft Gaussian mask. The B-factor was estimated to be −246.4 Å^2^. Local FSC calculation was performed using the Local Resolution tool in SPHIRE. (Supplementary figure 5e) and the electron density map was filtered according to its local resolution using the 3-D Local Filter tool in SPHIRE.

---

**Supplementary figure 1.**
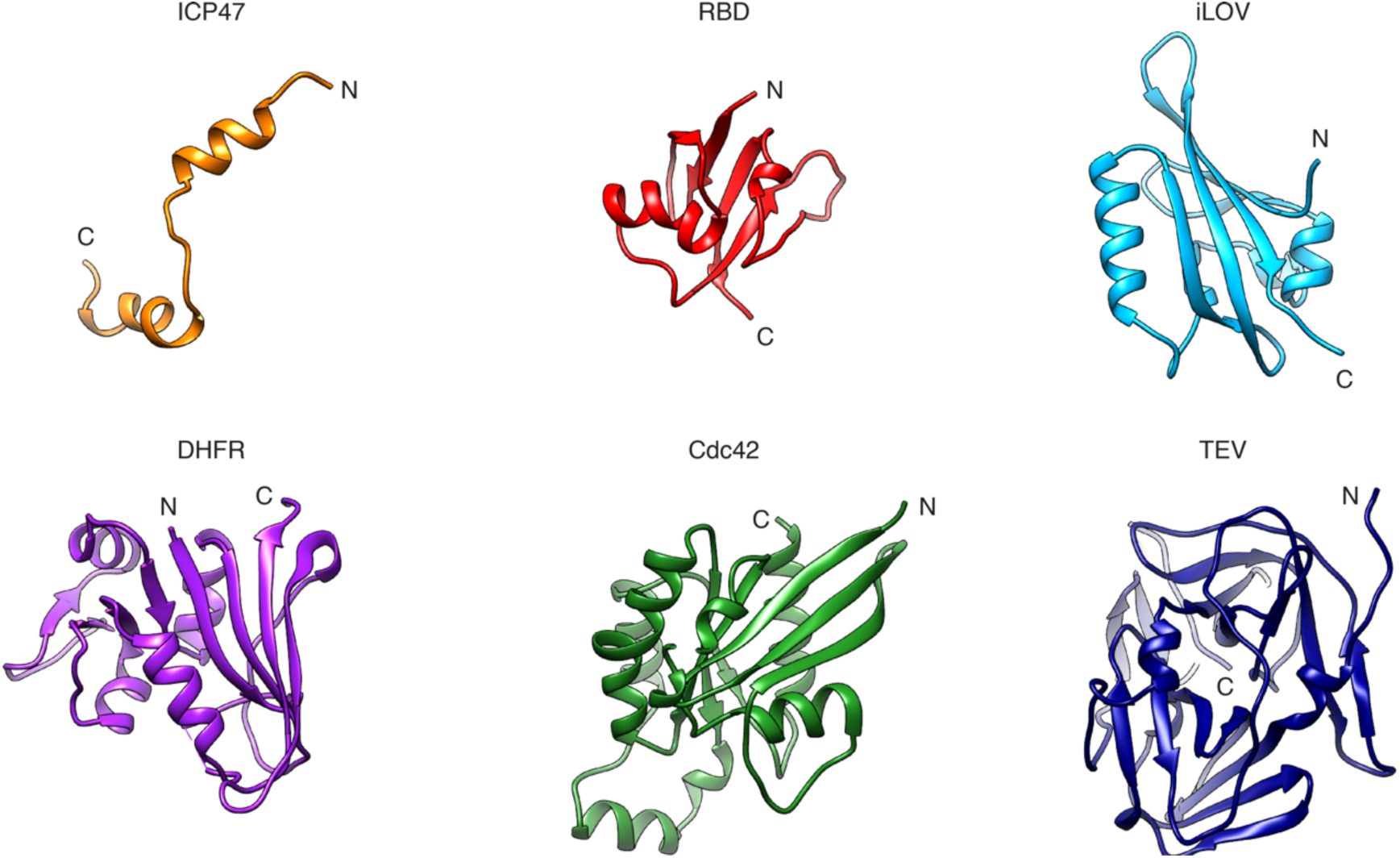
Structures of cargo proteins tested for translocation through Tc. Ribbon representation of the structures of the six cargo proteins tested in this study, with N- and C-termini indicated. ICP47: herpes simplex virus protein ICP47 (PDB ID 1QLO). RBD: Ras binding domain of human CRAF kinase (PDB ID 1RFA). iLOV: engineered LOV2 domain from *Arabidopsis thaliana* phototropin 2 (PDB ID 4EES). DHFR: *Escherichia coli* dihydrofolate reductase (PDB ID 6MT8). Cdc42: human small GTPase Cdc42 (PDB ID 4YC7). TEV: tobacco etch virus protease (PDB ID 1Q31). In the case of the NMR structure of the intrinsically disordered protein ICP47, only the N-terminal region (residues 1 – 34 out of 87) has been resolved. In case of Cdc42, one monomer of the homodimer is shown.

**Supplementary figure 2.**
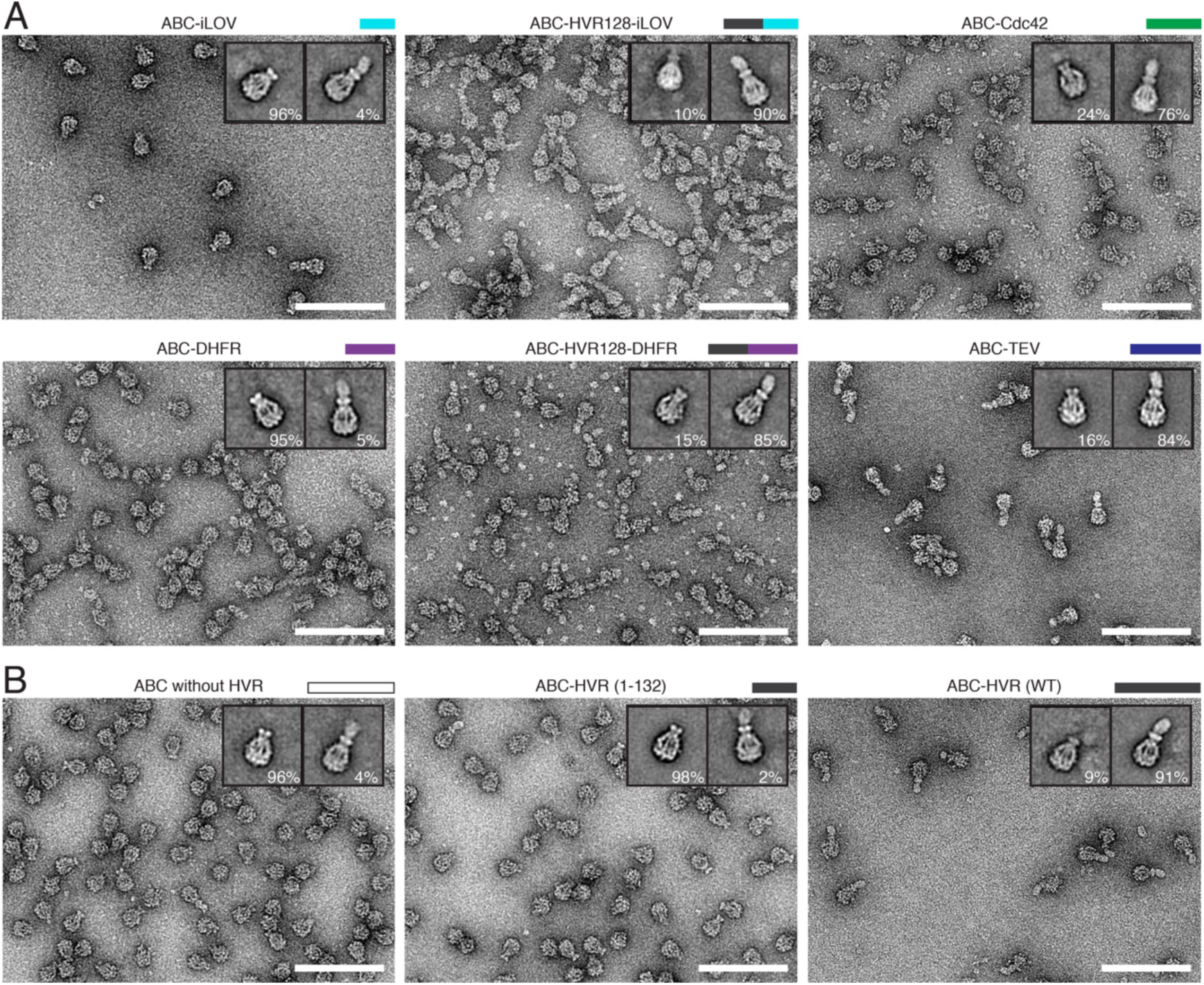
Formation of ABC holotoxin with different protein cargos in TcB-TcC. D. Negative stain electron micrographs of ABC after assembly of TcA with TcB-TcC cocoons containing different cargo proteins. Insets: 2D class averages of representative lone TcA pentamer and holotoxin, with the percentages of particles in the class averages shown below. TcB-TcC containing iLOV (13.2 kDa) and DHFR (18.4 kDa) as cargo do not form holotoxins with high affinity (left panels). Fusing the first 128 residues of TccC3HVR to these proteins results in respective cargo sizes of 26.4 and 31.6 kDa, which can efficiently form ABC holotoxin (middle panels). The cargo proteins Cdc42 (20.3 kDa) and TEV (28.1 kDa) form holotoxin with high affinity on their own (right panels).
E. Negative stain electron micrographs of ABC after assembly of TcA with an empty TcB-TcC cocoon (left), TcB-TcC with a truncated HVR (residues 1 – 132, middle) and ABC(WT) (right). The insets show 2D class averages like described in panel A. The empty cocoon and the cocoon with truncated HVR are inefficient at forming holotoxin. Scale bars: 100 nm.

**Supplementary figure 3.**
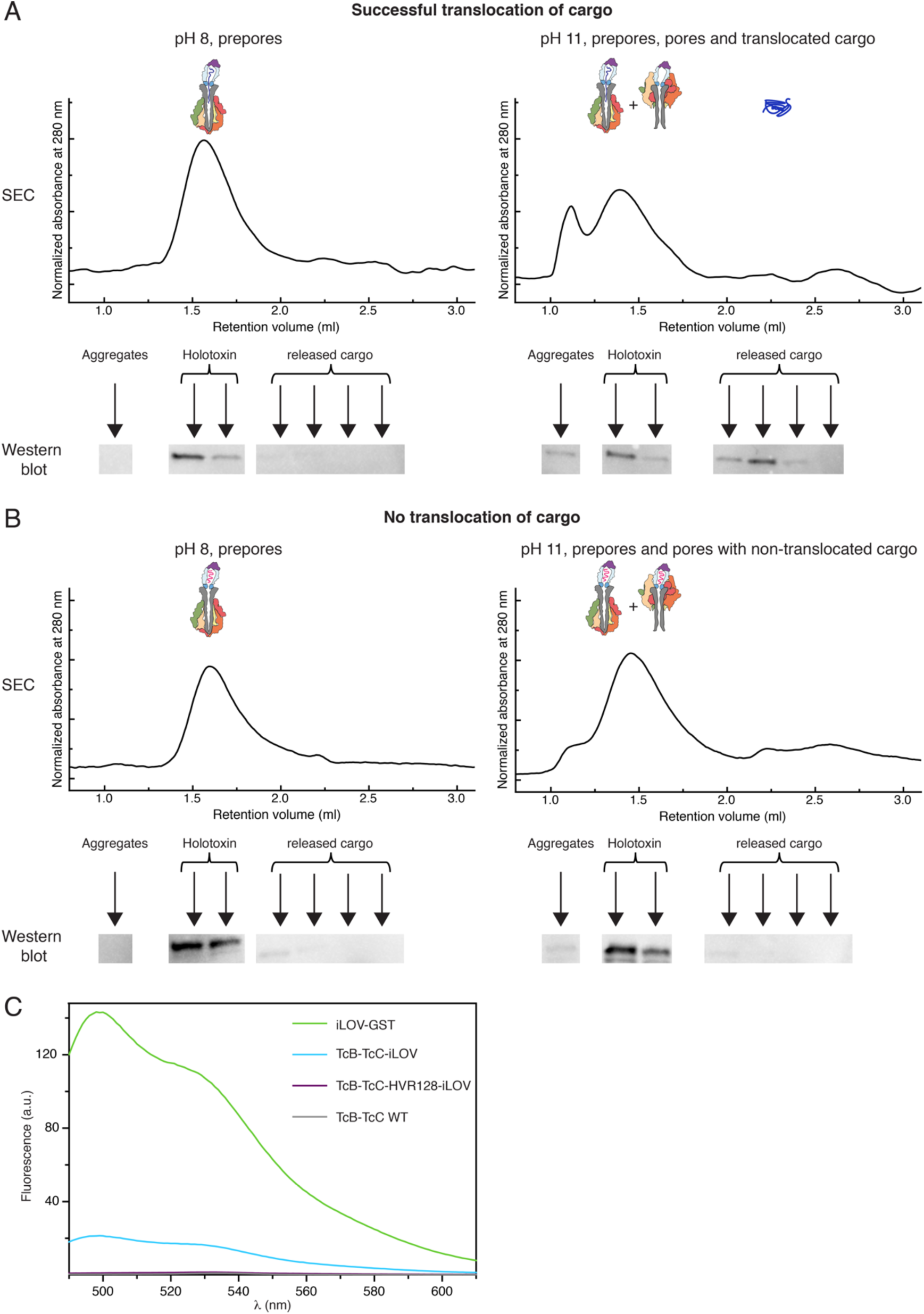
Scheme of the *in vitro* translocation experiment of cargo proteins and fluorescence spectroscopy of TcB-TcC-iLOV constructs. C. Successful translocation of the cargo. After incubating the holotoxin for 48 h at pH 8 (as a control where no prepore-to-pore transition occurs) and pH 11 (which induces prepore-to-pore transition), the samples are subjected to SEC on a Superose 6 Increase column. This separates aggregated holotoxin, holotoxin pores and residual prepores (lower retention volumes) from the translocated cargo (higher retention volumes). Subsequently, SEC fractions are analyzed for the presence of cargo via Western blot using antibodies raised either against the cargo proteins or against TccC3HVR (in case of TccC3HVR fusion constructs). At pH 8, the cargo is still in the cocoon and co-elutes with the holotoxin. At pH 11, the cargo has been released from the holotoxin that transited to the pore state, and therefore also appears at higher retention volumes in comparison to experiments done at pH 8.
D. No translocation of the cargo. After SEC of samples incubated at pH 11, Western blot analysis shows that the cargo co-elutes exclusively with the holotoxin and cannot be detected at higher retention volumes.
E. Fluorescence emission spectra of 500 nM TcB-TcC-iLOV and TcB-TcC-HVR128-iLOV in comparison with iLOV-GST and TcB-TcC WT. TcB-TcC-HVR128-iLOV is non-fluorescent, while TcB-TcC-iLOV shows ∼10% of the fluorescence of iLOV-GST.

**Supplementary figure 4.**
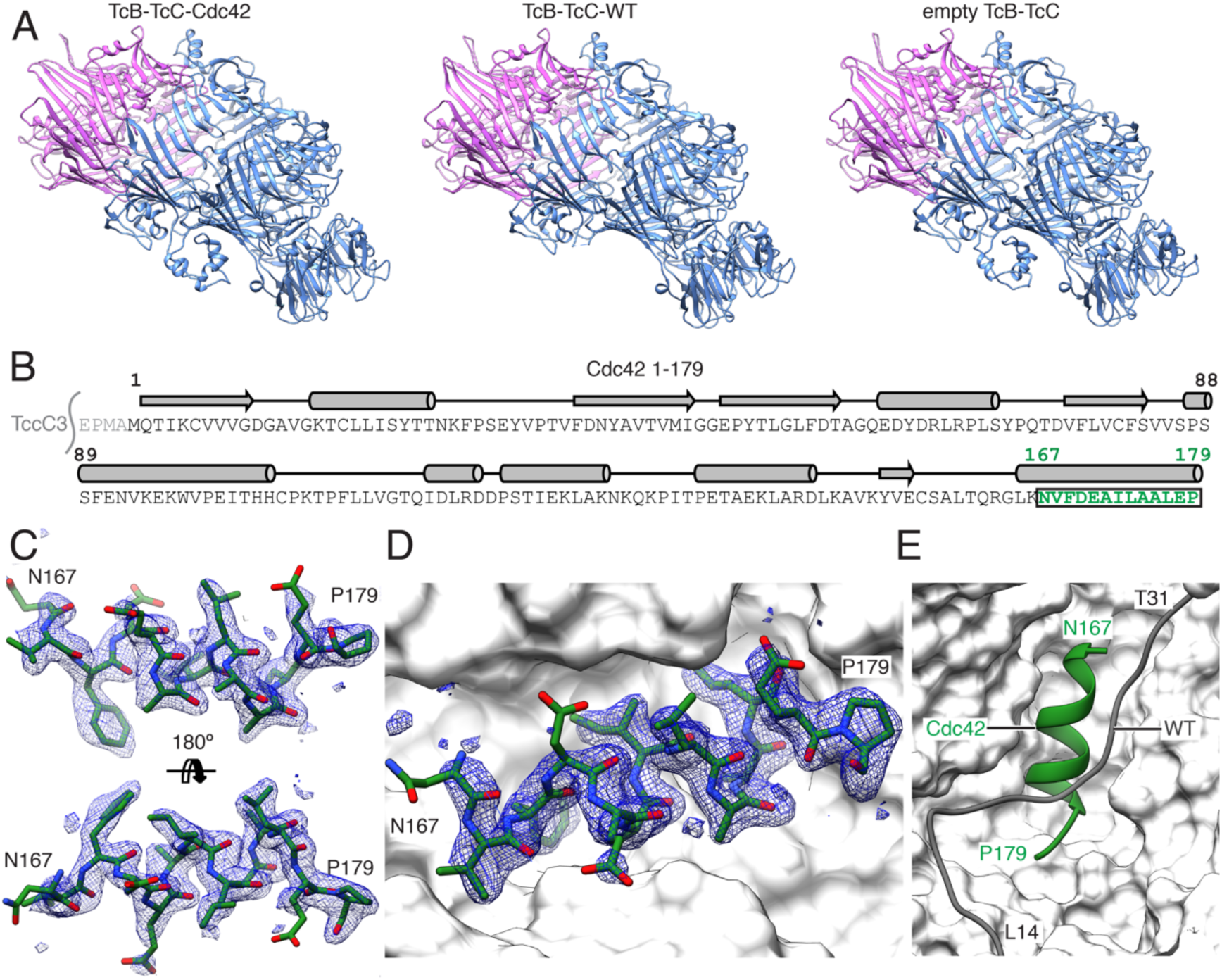
Crystallographic density of the Cdc42 C-terminus inside the TcB-TcC cocoon. A. Comparison of the TcB-TcC-Cdc42 (left, this work), TcB-TcC(WT) (middle, PDB ID 4O9X) and empty TcB-TcC (right, PDB ID 6H6G) structures.
B. Sequence of the Cdc42 cargo inside TcB-TcC. The first four residues after the TccC3 cleavage site (grey) are an N-terminal extension remaining from restriction cloning. The secondary structure according to PDB ID 4YC7 is indicated above the sequence. Residues 167 – 179 at the C-terminus (green) form the α-helix attached to the cocoon’s inner surface.
C. Model of the Cdc42 C-terminus in the crystallographic density map (blue mesh, contoured at 0.92 sigma). Two orientations are shown.
D. Model of the Cdc42 C-terminus in the crystallographic density map (blue mesh, contoured at 0.95 sigma) within the binding pocket of the cocoon (white surface).
E. Overlay of the binding pocket of TcB-TcC-Cdc42 including the Cdc42 C-terminus (green) with TcB-TcC(WT). The N-terminus of TcB-TcC(WT) (gray) occupies the space of Cdc42, and the corresponding N-terminal region of TcB-TcC-Cdc42 is not resolved.

**Supplementary figure 5.**
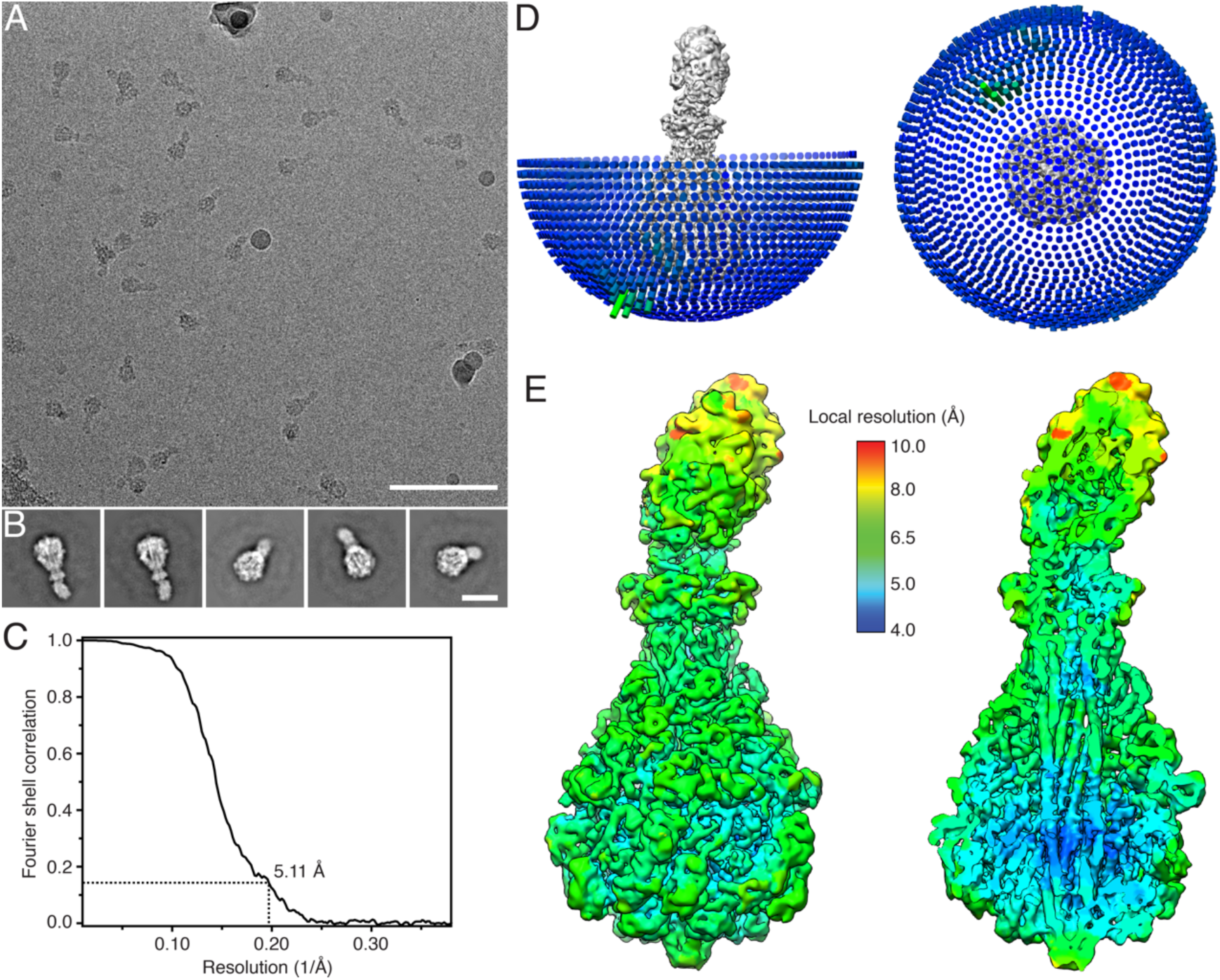
Cryo-EM of ABC-Cdc42. A. Representative digital electron micrograph of ABC-Cdc42 at 2.2 μm defocus and a total dose of 65 e^-^/Å^2^. The image was acquired with a Falcon II direct electron detector on a Cs corrected Titan Krios microscope. Scale bar: 100 nm.
B. Representative 2D class averages of ABC-Cdc42, showing side views and tilted views. Scale bar: 20 nm.
C. Fourier shell correlation (FSC) of the obtained cryo-EM map. The dashed line shows the 0.143 FSC cutoff criterion, with a resulting resolution of 5.11 Å.
D. Side and top view of the angular distribution plot from the final round of 3D refinement. Each cylinder composing the sphere represents a projection view at a particular angle, with length and color proportional to the number of particles at that projection angle. Longer and greener cylinders indicate more particles.
E. Side view (left) and cross section (right) of the ABC-Cdc42 cryo-EM density map colored according to the local resolution.

**Supplementary figure 6.**
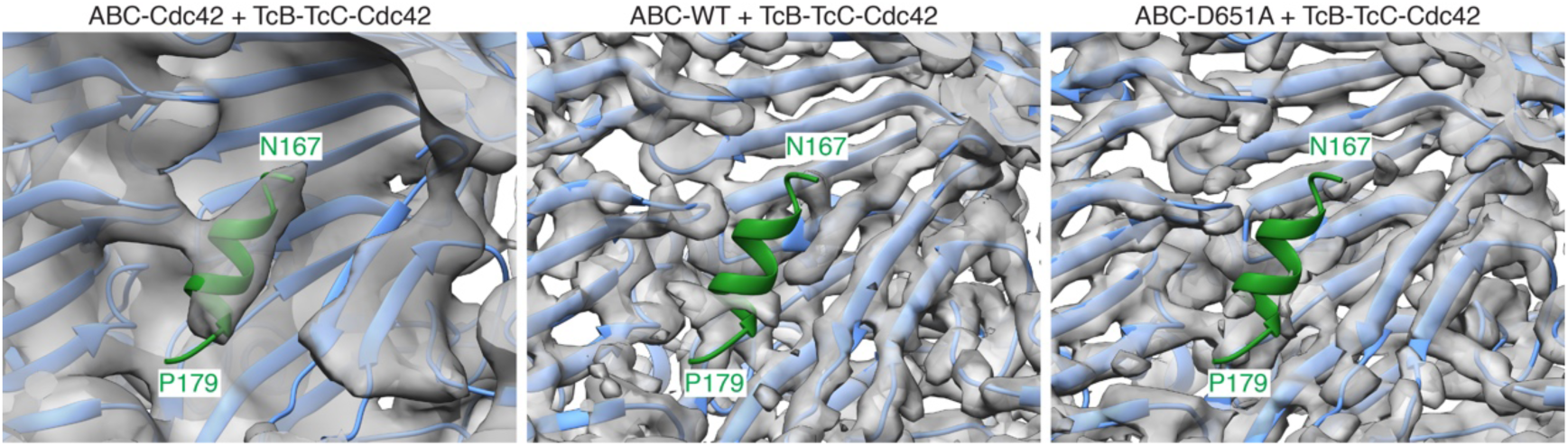
Visualization of density corresponding to the α-helix of Cdc42 in different ABC holotoxins. Overlay of the TcB-TcC-Cdc42 model with the cryo-EM maps of ABC-Cdc42 (left), ABC(WT) (EMDB 0149, center) and ABC-D651A (EMDB 0150, right). The C-terminal α-helix of Cdc42 is colored green, and the model of TcB is colored blue. Only ABC-Cdc42 shows density corresponding to a folded α-helix inside the cocoon.

**Supplementary table 1.** Data collection, processing and model refinement statistics of TcB-TcC-Cdc42. Values in brackets indicate the highest resolution shell. The table was generated with PHENIX.

| **Data collection** |  |
| --- | --- |
| Wavelength (Å) | 0.9796 |
| Space group | P2<sub>1</sub>2<sub>1</sub>2<sub>1</sub> |
| Cell dimensions |  |
| *a*, *b*, *c* (Å) | 96.35, 156.55, 179.46 |
| $\alpha$, $\beta$, $\gamma$ (°) | 90, 90, 90 |
| Resolution range (Å) | 48.17 - 2.0 (2.07 - 2.0) |
| Total reflections | 4174288 (319800) |
| Unique reflections | 182401 (18014) |
| Multiplicity | 22.9 (17.8) |
| *R*<sub>merge</sub> | 0.1855 (2.02) |
| *R*-*meas* | 0.1897 (2.08) |
| R-pim | 0.0393 (0.4803) |
| *I* / σ*I* | 13.81 (1.46) |
| Completeness (%) | 99.50 (99.37) |
| **Refinement** |  |
| No. reflections | 182143 (18014) |
| No. reflections R-free | 9111 (901) |
| *R*<sub>work</sub> / *R*<sub>free</sub> (%) | 21.20 (28.70)/ 25.03 (32.02) |
| No. atoms | 18289 |
| Protein | 17004 |
| Ligand/ion | 3 |
| Water | 1282 |
| *B*-factors |  |
| Protein | 41.86 |
| Ligand/ion | 35.60 |
| Water | 42.20 |
| R.m.s. deviations |  |
| Bond lengths (Å) | 0.005 |
| Bond angles (°) | 0.78 |
| Ramachandran favoured (%) | 97.08 |
| Ramachandran allowed (%) | 2.78 |
| Ramachandran outliers (%) | 0.14 |
| Rotamer outliers (%) | 0.38 |
| Clashscore | 6.08 |

**Supplementary table 2.** Cryo-EM data processing statistics of ABC-Cdc42.

| **Data collection and processing** |  |
| --- | --- |
| Microscope | Titan Krios (Cs corrected, XFEG) |
| Camera | FEI Falcon II |
| Magnification | 59,000x |
| Voltage (kV) | 300 |
| Number of frames | 24 |
| Exposure time (s) | 1.5 |
| Total electron exposure ($e^-/\text{\AA}^2$) | 65 |
| Defocus range ($\mu\text{m}$) | 1.0 – 3.2 |
| Pixel size ($\text{\AA}$) | 1.14 |
| Symmetry imposed | C1 |
| Initial particles (no.) | 99,980 |
| Final particles (no.) | 56,665 |
| Map resolution ($\text{\AA}$) | 5.1 |
| FSC threshold | 0.143 |
| Map sharpening $B$ factor ($\text{\AA}^2$) | -246.4 |

